# Endothelial and neuronal engagement by AAV-BR1 alleviates neurological symptoms and cholesterol deposition in a mouse model of Niemann-Pick type C2

**DOI:** 10.1101/2024.04.15.589486

**Authors:** Charlotte Laurfelt Munch Rasmussen, Christian Würtz Heegaard, Maj Schneider Thomsen, Eva Hede, Bartosz Laczek, Jakob Körbelin, Louiza Bohn Thomsen, Markus Schwaninger, Torben Moos, Annette Burkhart

## Abstract

**Background:** Patients with the genetic disorder Niemann-Pick type C2 disease (NP-C2) suffer from lysosomal accumulation of cholesterol causing both systemic and severe neurological symptoms. In a murine NP-C2 model, otherwise successful intravenous Niemann-Pick C2 protein (NPC2) replacement therapy fails to alleviate progressive neurodegeneration as infused NPC2 is unable to cross the blood-brain barrier (BBB). Genetic modification of brain endothelial cells (BECs) is thought to enable secretion of recombinant proteins thereby overcoming the restrictions of the BBB. We hypothesized that BBB-directed gene therapy using the AAV-BR1-NPC2 vector would transduce both BECs and neurons in a mouse model of NP-C2 (*Npc2*-/-).

**Methods:** Six weeks old *Npc2*-/- mice were intravenously injected with the AAV-BR1-NPC2 vector. Post-mortem analyses included gene expression analyses, determination of NPC2 transduction in the CNS, and co-detection of cholesterol with NPC2 in neurons.

**Results:** The vector exerted tropism for BECs and neurons resulting in a widespread NPC2 distribution in the brain with a concomitant reduction of cholesterol in adjacent neurons, presumably not transduced by the vector.

**Conclusion:** The data suggests cross-correcting gene therapy to the brain via delivery of NPC2 from BECs and neurons.

## Letter Manuscript

Niemann-Pick (NP) type C disease (NP-C) is an autosomal recessive lysosomal storage disease caused by mutations in the *Npc1* or *Npc2* genes leading to neurodegeneration (1). Loss of function of either of the resulting NP-C proteins results in lysosomal accumulation of cholesterol leading to human diseases NP-C1 and NP-C2 (2). The glycoprotein NPC2 is found in lysosomes and secretory fluids, and secreted NPC2 is endocytosed via the ubiquitous mannose-6-phosphate receptor (M6PR) pathway (3). Transport of NPC2 from blood to the brain is restricted because of the blood-brain barrier (BBB) formed by brain endothelial cells (BECs) and because M6PR expression by BECs declines after birth (4).

NP-C2 progresses with fatal outcomes, and most patients die between 10 and 25 years of age. Curative treatment of NP-C2 is unavailable, and the development of new treatments is highly warranted (5). Genetic modification using a capsid-modified adeno-associated virus vector (AAV-BR1), known for its robust tropism for BECs, enables recombinant NPC2 secretion *in vitro* (3), suggesting that NPC2 released from BECs also might access the intact brain. The therapeutic potential of AAV-BR1 was recently verified in mouse models of neurodegeneration, i.e. Incontinentia pigmenti, Sandhoff disease, and Allan-Herndon-Dudley syndrome (6–8), with successful treatment enabled by induced gene expression in BECs. We recently showed that systemic administration of AAV-BR1 encoding NPC2 in healthy mice somewhat surprisingly transduced both BECs and neurons (3), suggesting that AAV-BR1 passes the BBB and enables neuronal transduction. Accordingly, we hypothesized that BBB-directed gene therapy using the AAV-BR1-NPC2 vector would transduce both BECs and neurons in a mouse model of NP-C2 (*Npc2*-/-).

The present study demonstrates that intravenous (IV) injection of the AAV-BR1-NPC2 vector in NPC2-deficient mice aged six weeks evokes the distribution of NPC2 in BECs and neurons and lowers neuronal cholesterol. *Npc2*-/-mice exhibit normal growth patterns similar to wildtype (WT) littermates until 6 weeks of age at which a single IV dose of 1.6 x 10^11^ vg of the AAV-BR1-NPC2 vector was administered to *Npc2*-/- mice. At 12 weeks, untreated and AAV-BR1-NPC2-treated *Npc2*-/- mice revealed growth retardation, similar to that reported previously (10).

In the *Npc2*-/- mice, the AAV-BR1-NPC2 vector accumulated in brain, lung, and lightly in spleen (Fig.1A). We subsequently compared *Npc2* gene expression in these organs with levels found in untreated *Npc2*-/- and WT littermates using qPCR (Supplementary Table S1). High accumulation of AAV-BR1-NPC2 vector in the brain increased the *Npc2* gene expression to a level not significantly different from that of *Npc2+/+* mice. Only a slight increase in *Npc2* gene expression was seen in the lung after AAV-BR1 treatment despite high vector accumulation (Fig.1A). *Npc2* gene expression was absent from the spleen (data not shown), suggesting unspecific uptake of the AAV-BR1 vector. Cerebral distribution of NPC2 after transduction with the AAV-BR1-NPC2 vector was evaluated using immunohistochemistry. Injecting the vector revealed NPC2-positive neurons, especially in the cerebral cortex and hippocampus (Fig.1A), and dispersed in the striatum, thalamus, hypothalamus, mesencephalon, pons, medulla oblongata, and cerebellum (not shown). Neurons were intensively stained with a weaker, yet consistent, staining of several brain capillaries. NPC2-immunoreactive cells with morphology corresponding to astrocytes, microglia, or oligodendrocytes were not observed. NPC2 protein was undetectable in untreated *Npc2*-/- mice. There were variations in transduction efficiency among the treated mice, which correlated with an increased number of NPC2-positive cells, and a higher therapeutic potential.

**Legend to Figure 1.**
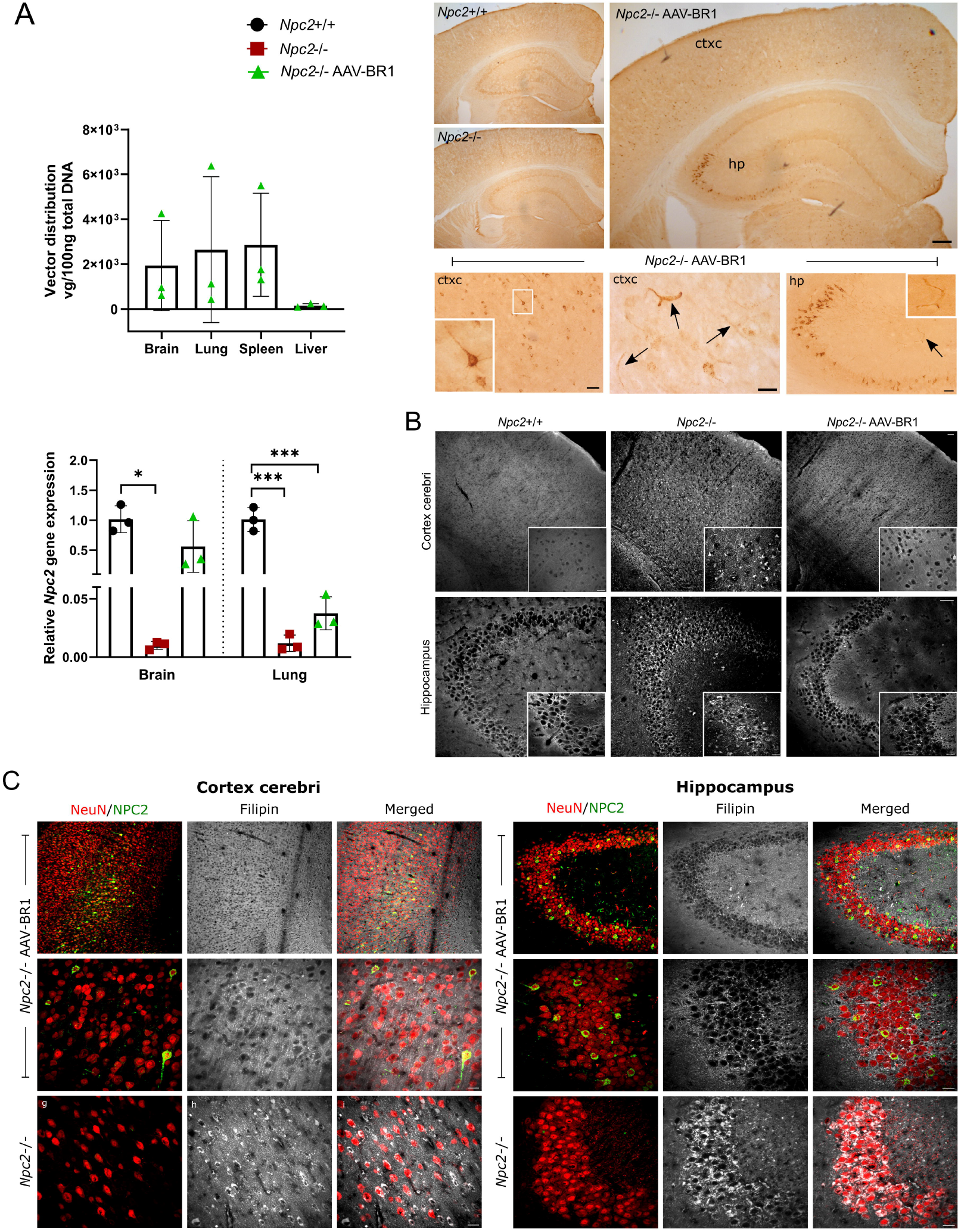
**A** **Upper left**, Biodistribution of AAV-BR1-NPC2 (viral genomes (vg)) in brain, lung, liver, and spleen, analyzed by quantitative qPCR at 12 weeks (n=3). **Lower left**, Relative *Npc2* gene expression in the brain and lung analyzed by RT-qPCR in *Npc2*+/+ (n=3, black), *Npc2*- /- (n=3, red), and AAV-BR1-NPC2-treated *Npc2*-/- mice (n=3, green). *Npc2* expression is significantly lower in the brains of untreated *Npc2*-/- mice. *Npc2* expression in the lungs of treated and untreated *Npc2*-/- mice is significantly lower compared to wild-type littermates (Mean ± SD). Data are analyzed with a one-way ANOVA (F_Cerebrum_[2,6] = 9.64, *p* = 0.013, F_Lung_[2,6] = 74.52, *p* < 0.0001) with Tukey’s multiple comparisons test (* *p* = 0.011, *** *p* ≤ 0.0001). **Upper right**, Immunohistochemistry reveals NPC2-positive cells in the brain of *Npc2*-/- mice after AAV-BR1-NPC2 gene therapy, particularly prominent in neurons of cortex cerebri (ctxc) and hippocampus (hp) (CA3 region). NPC2 is also seen in brain capillaries (arrows). Scale bars = 50 µm (ctxc and hp), 25 µm (lower row). **B**. Cholesterol accumulation (visualized as white elements in filipin staining) is virtually absent in cortex cerebri of *Npc2*+/+ mice, clearly present in *Npc2*-/- mice, and reduced when treated with AAV-BR1-NPC2. Scale bars = 50 µm and 20 µm (white boxes). **C**. Neurons (visualized with neuronal nuclei antigen (NeuN) immunolabeling (red)) also exhibit NPC2 immunoreactivity (green, arrows)) in cortex cerebri and hippocampus in AAV-BR1-NPC2-treated *Npc2*-/- mice. Of note, AAV-BR1-NPC2-treated *Npc2*-/- mice are virtually devoid of filipin labeling, clearly contrasted by the robust appearance in untreated *Npc2*-/- mice. In *Npc2*-/- mice, treatment reduction of cholesterol accumulation is not limited to NPC2 expressing cells, but also present in neighboring cells suggesting uptake of secreted NPC2 equivalent to cross-correction. Scale bar = 25µm.

Cholesterol accumulation in the cortex cerebri and hippocampus visualized using filipin, a fluorescent antibiotic binding specifically to unesterified cholesterol, was undetectable in WT mice while present in *Npc2*-/- mice with reduction after AAV-BR1-NPC2-treatment (Fig.1B). As reduced cholesterol appeared in areas with many NPC2-containing neurons after AAV-BR1-NPC2-treatment, we aimed for co-detection of filipin and NPC2 focusing on cortex cerebri and hippocampus (Fig.1C). In cortex cerebri of AAV-BR1-NPC2-treated *Npc2*-/- mice, superimposing immunolabeled images with filipin staining evidenced that NPC2 neurons contained cholesterol but to a much lesser extent than seen in untreated *Npc2*-/- mice (Fig.1C, left panel). Similarly, the high number of NPC2-positive neurons found in the CA3 region of the hippocampus in AAV-BR1-NPC2-treated *Npc2*-/- mice was almost completely reversed concerning cholesterol accumulation (Fig.1C, right panel). The number of NPC2-labeled neurons in AAV-BR1-NPC2-treated *Npc2*-/- mice was lower than that of cholesterol-negative cells in both cerebral cortex and hippocampus, which suggests that secreted NPC2 from AAV-BR1-NPC2-transduced neurons alleviates cholesterol accumulation in non-transduced neighboring neurons corresponding to neuronal cross-correction (6).

Collectively, our data showed that BBB-directed gene therapy using the AAV-BR1 vector led to the widespread appearance of NPC2 in BECs and neurons, which was accompanied by reduction in neuronal cholesterol in the cerebral cortex and hippocampus.

The integrity of the BBB is often compromised during pathological conditions, but this is not expected in the *Npc2*-/- mice. NPC2 replacement therapy in a mouse model of NP-C2 similar to that of the present study mended visceral cholesterol storage but failed to improve neurological symptoms after repeated intravenous injections with NPC2, stating the incapability of NPC2 to cross the BBB (c.f. 3). This was emphasized previously where improvement in the BBB integrity was observed after intravenous injections of the AAV-BR1-vector in mice suffering from Incontinentia pigmenti (6,8). There is, therefore, no reason to believe that the AAV-BR1 vector passes the BBB and transduces neurons due to compromised barrier integrity in the *Npc2*-/- mice. However, the mechanism of this BBB passage remains unknown. Loss of BBB integrity seen in pathological conditions was not observed in *Npc2*-/- mice (9). The predominant appearance of NPC2-positive cells was in the hippocampus consistent with studies using healthy mice (9). Compared to evolutionary higher regions, the hippocampal vessels differ by lesser neurovascular coupling between vascular networks and functions of BECs and pericytes, which may facilitate viral vector binding and capillary migration.

The distribution pattern of NPC2 in the neurons was similar to our previous study investigating the AAV-BR1-NPC2 in healthy BALB/cJRj mice (9). Here the AAV-BR1 vector was designed to produce two separate proteins; enhanced green fluorescent protein (eGFP) and NPC2, where the eGFP accumulated intracellularly as an indicator of cellular transduction. However, also neuronal cells expressed eGFP, indicating that some of the systemically injected AAV-BR1 undergoes transport across the BBB leading to neuronal transduction (9). In our previous study in healthy mice (9), we further examined the possibility of AAV-BR1 vector transport across the blood-cerebrospinal fluid barrier (BCSFB) but found no evidence to support this. Furthermore, it seems unlikely that the widespread distribution of NPC2 throughout the brain, including the cerebral cortex, could be achieved through BCSFB transport.

When examining the correlation between the distribution of NPC2 and cholesterol storage in both the cerebral cortex and hippocampus, a clear reduction in cholesterol accumulation was seen in NPC2-positive neurons but also neighboring NPC2-negative cells, emphasizing the possibility of cross-correction after BBB-directed gene therapy. The heterogeneous, albeit substantial, neuronal presence of NPC2, might be a reflection of cation-independent M6PR mediated uptake of the mannose 6-phosphate tagged NPC2 (3), as cation-independent M6PR also distributes to neurons quite heterogeneously in the brain with a predominantly high occurrence in deep layers of the cortex (e.g., pyramidal neurons in layer V), neurons of the hippocampus, striatum, selected nuclei in the thalamus, Purkinje cells of the cerebellum, the deep cerebellar nuclei, red nucleus, pontine nucleus, and motor neurons of the brainstem (c.f. 3), corresponding well to the areas where NPC2-positive neurons are observed.

In conclusion, BBB-directed gene therapy using the AAV-BR1 vector corresponded with widespread neuronal expression of NPC2 and reduction of cholesterol accumulation. Cortex cerebri and hippocampus are severely affected by neurodegeneration in mouse models of NP-C and human NP-C and display intraneuronal lipid deposition, especially in large pyramidal neurons. The high transduction in cortex cerebri and hippocampus reduced cholesterol accumulation in both genetically modified (NPC2+) and non-transduced (NPC2-) neurons suggesting cross-correcting gene therapy via secretion of NPC2 from BECs and NPC2+ neurons.

## Acknowledgments

The authors thank laboratory technicians Merete Fredsgaard, Hanne Krone Nielsen, Ditte Bech Laursen, and Louise Hvilshøj Madsen, Aalborg University, Denmark, and animal technicians Karina Lassen Holm and Dorte Hermansen, Aarhus University, Denmark, for excellent technical assistance during the study. Associate Professor Anders Olsen and Helene Halkjær Jensen, Department of Chemistry and Bioscience, Aalborg University are acknowledged for assistance with the use of the Olympus IX83 inverted microscope equipped with Yokogawa confocal CSU-W1 spinning disk.

## Abbreviations

AAV-BR1: adeno-associated virus vector
BR-1; BBB: blood-brain barrier
BCSFB: blood-cerebrospinal fluid barrier
BECs: brain endothelial cells
IV: intravenous
NP: Niemann-Pick
NP-C: Niemann-Pick type C disease
WT: wildtype

