## Supplementary material for "Endothelial and neuronal engagement by AAV-BR1 alleviates neurological symptoms and cholesterol deposition in a mouse model of Niemann-Pick type C2": Suppl Material

**Materials and methods
Production of recombinant viral vectors**

The plasmid pAAV-CAG-NPC2 was generated following the procedure described earlier (1). The recombinant AAV-BR1-NPC2 vector used in the present study was produced by co-infection of Sf9 cells using the baculovirus expression system as previously described (2). In short, Sf9 cells were infected with two different recombinant baculoviruses; one carrying the AAV2 *rep* gene and a brain-endothelial cell-specific AAV2-BR1 *cap* gene, and one containing the CAG promoter and the mouse *Npc2* gene flanked by inverted terminal repeats from AAV2. Four days after the co-infection, the viral particles were harvested from the Sf9 cells by repeated freeze/thaw cycles and digested with 50 U/mL Benzonase Nuclease to remove unpackaged DNA. The viral vectors were purified using iodixanol density-gradient ultracentrifugation and extracted from the 40 % iodixanol layer. Genomic titers were determined by quantitative real-time PCR (qPCR) using CAG-specific primers (Table S1). The SYBR Green-based FastStart Essential DNA Green Master (Merck KGaA, #06402712001 Roche) with the Light Cycler 96 System (Roche) was used to determine the vector copy numbers. The reactions were run with an initial denaturation for 10 min at 95°C followed by 40 cycles of denaturation for 30 s at 95°C, then annealing for 30 s at 67°C, and extension for 30 s at 72°C, followed by a melt curve analysis (60–97°C, 0.1°C/s).

**Animal study approval and reporting**

The animal studies were performed according to the Danish Animal Experimentation Act (BEK no. 2028 of 14/12/2020) and the European directive (2010/63/EU) and carried out by licensed staff. The Danish Animal Experiments Inspectorate under the Ministry of Food, Fisheries, and Agriculture has approved all animal experiments and breeding of NPC2-deficient mice (license no. 2018-15-0201-01467 and 2019-15-0202-00056).

**Animals**

The 129P2/OlaHsd-*Npc2^Gt(LST105)BygNya^* mouse strain (4) was rederived using *in vitro* fertilization, and the strain established on a BALB/cJRj background (supplied by Janvier Labs, Le Genest‐Saint‐Isle, France). Heterozygous *Npc2*+/- mice were mated to obtain offspring homozygous for the mutation (*Npc2*-/-) and wild-type (WT) control littermates (*Npc2+/+*). The breeding was established using continuous trio-breeding (two *Npc2*+/- female mice were housed with one *Npc2*+/- male mouse) (5). The housing and breeding were carried out at the animal facility at Aarhus University, Aarhus, Denmark, under specific pathogen-free conditions. Health monitoring was followed according to FELASA recommendations (6), and the mice were free of all pathogens listed. Both female and male mice were included in the study. The mice were group-housed with up to five mice per cage in standard IVC cages (GM500, Tecniplast) under controlled conditions (ambient temperature 20-24°C, 55 ± 10% humidity, and a 12-hour light/dark cycle with the light on at 6:00 am). They were provided with environmental enrichment consisting of Tapvei bedding material, sizzle nesting material, aspen bricks, biodegradable cardboard houses, tunnels, and peanuts as food enrichment. The cages were changed once a week. Offspring were weaned at 21 days of age and separated according to sex. Mice were provided with a standard chow diet (Altromin #1324, Brogaarden, Lynge, Denmark) and reverse osmosis water *ad libitum*.

**Genotyping**

Genotyping of mice was performed on postnatal days 10-14 based on ear punching (5) which was also used for individual identification. DNA from ear punches was purified using the Quick-DNA mini prep plus kit (Zymo Research # D4068) according to the manufacturer’s protocol. DNA quantity and purity were determined with a DeNovix Spectrophotometer DS-11. The *Npc2*-/- genotype was determined by qPCRs using Maxima SYBR Green Master Mix, with ROX as a reference (Thermo Fisher Scientific, #K0223) and two sets of primers (Table S1). All samples were analyzed in duplicates. The qPCR reaction was performed using the QuantStudio 6 Flex Real-Time PCR System (Thermo Fisher Scientific). In short, the qPCR reaction was initiated with denaturation of the samples at 95 °C for 10 minutes followed by 40 cycles of denaturation at 95 °C for 15 seconds, annealing at 60 °C for 30 seconds, and extension at 72°C for 30 seconds. Finally, a melt curve analysis was performed (60 °C to 95 °C with 0.05 °C/s).

***In vivo* study design**

A total of 30 mice were used in the study. Offspring from the trio breeding of heterozygote *Npc2*+/- were divided into three groups (n=10/group). The *Npc2*-/- mice were randomly allocated to their respective group (treated vs. untreated) using a computer-based random order generator (7). The study director was blinded to treatment status during the entire study period and blinded to genotype and treatment group during the histopathological evaluation. In the first and second groups, six weeks old *Npc2*-/- mice were intravenously injected with sterile phosphate-buffered saline (PBS) or AAV-BR1-NPC2 vectors diluted in PBS (dose at 1.6x10^11^ viral vectors/mouse) in the tail vein. Both groups were injected with the same volume (200 µl). The vector dose and route of administration are based on previous studies (1,2). The third group included age-matched untreated wild-type (WT) littermates used as controls. The experimental unit is the individual mouse, and since only 1 in 7 mice is born with the *Npc2*-/- genotype (5), the study was conducted in seven cohorts. Whenever possible, all experimental groups were represented in each of the cages. During the study period, the mice were monitored daily for their clinical condition. The body weight was measured weekly, but from 60 days of age until the end of the study (day 84), the body weight was measured twice a week. At 12 weeks of age, before reaching the end stage of the disease, the study was terminated.

**Human endpoints**

Based on knowledge of disease progression and manifestations from a previous study using similar *Npc2-/-* mice (4), the mice were euthanized at 12 weeks of age (six weeks post-injection) or whenever humane endpoints were reached. The humane endpoints were as follows: inability to drink or eat, dehydration, severe tremors, and ataxia resulting in repeated falling or reluctance to move, weighing 20 % less compared to WT mice of identical age and sex, and penile prolapse.

**Tissue collection and processing**

At 12 weeks of age, the mice were deeply anesthetized with 5 % isoflurane (1L/min O_2_), and once reflexes were absent, the thoracic cavity was opened, and the animals transcardially perfused with a manual injection of 20 ml cold PBS. Brain, liver, spleen, and lung tissue were harvested from every animal (n=10) and weighed using a precision scale. Afterward, the organs were divided in two; one half was snap-frozen on dry ice and stored at -70°C for biochemical analysis, and the other half was post-fixated in 4 % PFA overnight at 4°C. Post-fixated organs were subsequently thoroughly washed in potassium-containing phosphate-buffered saline (PPBS) and stored at 4°C in PPBS with 0.1 % sodium azide.

**Quantification of viral genomes in tissue**

Approximately 10-20 mg tissue from the cerebrum, lung, liver, and spleen was homogenized in RNeasy Lysis Buffer with mercaptoethanol, and the DNA and RNA (gene expression analysis, see next section) were purified using the AllPrep DNA/RNA Mini Kit (Qiagen, #80204) according to the manufacturer’s protocol. The purity and quantity of the DNA and RNA were assessed by a spectrophotometer DS-11 (DeNovix). To analyze the organ distribution of the AAV-BR1-NPC2 vector, absolute qPCR was used to quantify the number of vg. See the section “Production of recombinant viral vectors” for the qPCR protocol. 100 ng DNA was used for each sample, which was run in triplicates. The quantification of vg was generated using a standard curve based on plasmid DNA encoding the CAG promoter as previously described (2). Data are reported as vg/100ng total DNA.

***Npc2* gene expression analysis**

RNA, extracted from the cerebrum, liver, lung, and spleen as described above, was treated with DNase I enzyme (Thermo Fischer Scientific, #EN0521) to remove genomic DNA contamination. 100 ng DNA-free RNA was used as the template for the cDNA synthesis, which was performed as previously described (2). The *Npc2* gene expression was analyzed using a probe-based multiplex RT-qPCR with Taqman Fast Advanced Master Mix (Thermo Fischer Scientific, #4444556), FAM conjugated mouse *Npc2* primer-probe mix (Thermo Fischer Scientific, #4331182, assay ID: Mm00499230_m1), and VIC-conjugated mouse hypoxanthine phosphoribosyltransferase (*Hprt1)* primer-probe mix (Thermo Fischer Scientific, #4448490, assay ID: Hs02800695_m1). *Hprt1* was used as the reference gene. 5 ng cDNA was used in the PCR reaction (For qPCR specifications see (2)). Finally, the relative gene expression ratios of *Npc2* were calculated using the delta-delta threshold cycle (Ct) method with the WT control mice as the reference sample.

**Morphological investigations**

40 µm cryosections of brain tissue were prepared as previously described (2). Free-floating brain sections were incubated in 3 % porcine serum with 0.3 % Triton X-100 diluted in 0.1 M PPBS (blocking buffer) for 30 minutes at room temperature to block unspecific binding and permeabilize cell membranes. Then the sections were incubated overnight at 4°C with the following primary antibodies diluted in blocking buffer: Neuronal nuclei antigen (NeuN) (Chemicon, #MAB377) (1:500) and anti-bovine NPC2 (Immuno-affinity purified antibody derived from anti-NPC2 IgG positive rabbit serum produced by subcutaneous injections of native bovine NPC2 (4) (1:2,000)). After three washes in blocking buffer diluted 1:50 in PPBS (washing buffer), most sections were incubated with biotinylated goat anti-rabbit IgG (Vector, #BA-1000) diluted 1:200 in blocking buffer for one hour. Sections were washed twice in washing buffer and once in PPBS. For visualization, the sections were incubated in an Avidin-Biotin Complex (ABComplex)-system (VECTASTAIN® Elite ABC-HRP Kit, Vector laboratories, #PK6100) diluted in PPBS and finally in 3,3′-diaminobenzidine tetrahydrochloride (DAB). The sections were mounted with Pertex. Section double stained for NeuN and NPC2 were visualized using goat anti-mouse Alexa 594 (Invitrogen #A11032) (1:200), and biotinylated goat anti-rabbit (1:200). For tyramide enhancement of NPC2 (2), ABComplex Vectastain were added for 30 minutes and washed in PPBS. The Tyramide Signal Amplification (TSA) Biotin kit (AKOYA Biosciences, #NEL700A001KT) was added for five minutes, and the sections were washed in PPBS and incubated with the ABComplex Vectastain for 30 minutes. The sections were again washed in PPBS and visualized using an anti-streptavidin Alexa 488 (Invitrogen, #S32354) (1:200) antibody for one hour. These sections were subsequently stained with Filipin, as explained below.

Cholesterol staining on brain sections was performed using filipin complex III (Sigma-Aldrich, #F4767) modified from (8). Filipin is a fluorescent antibiotic that specifically binds unesterified cholesterol (9). Filipin stock solution in dimethyl sulfoxide (1mg/ml) was diluted in PBS for a final working solution of 50 µg/ml. The free-floating brain sections were washed twice in PPBS, followed by two times washing in 0.02 % saponin in PBS. Sections were then incubated in a quenching solution consisting of 1.5 mg/ml glycine, 1 % bovine serum albumin, and 0.02 % saponin/PBS at room temperature for 30 minutes. Finally, the sections were incubated in Filipin working solution for three hours in the dark under agitation, washed twice in 0.02 % saponin in PBS, and mounted with DAKO fluorescent mounting media (#S3023). Images of DAB stains were acquired using an Axioplan 2 microscope equipped with an Axiocam MRc camera (Carl Zeiss). Filipin stain were imaged using an Olympus IX83 inverted microscope with a Yokogawa confocal CSU-W1 spinning disk unit equipped with a Hamamatsu ORCAFlash 4.0 v3 grayscale camera using an Olympus UPlanSApo 60x/1.35na oil objective. All images were generated using the same acquisition settings, analyzed with ImageJ software (10), and adjusted for brightness and contrast.

**Statistical analysis**

The sample size was calculated by power analysis using the G*Power software (version 3.1.9.2). The sample size was calculated using an effect size of 1.5, which was based on previous studies evaluating the effect of gene therapy in NP-C mice (8). The power and significance level of the experiment was set to 80 % and 0.05, respectively. A loss of 10 % of the mice was expected based on the humane endpoints, and therefore 10 mice were included in each group. All data sets were analyzed by the D’Agostino-Pearson test to verify if the data followed a Gaussian distribution and tested for equal variances by the Brown-Forsythe test. If the data sets passed these tests, a one-way ANOVA with Tukey’s post hoc analysis was applied. Otherwise, the data sets were log-transformed. If the data sets still did not pass the parametric criteria, the Kruskal-Wallis test with Dunn’s multiple comparisons was applied. The details of the specific statistical analysis used are reported in the figure legends. All statistical analyses were two-sided. A p-value of ≤ 0.05 was considered statistically significant. Data are reported as mean ± standard deviation (SD) if not stated otherwise.

**Supplementary Table**

**Table S1. Overview of the primers used in the study.**

| **Gene** | **Forward** | **Reverse** |
| --- | --- | --- |

| *CAG* | AACGCCAATAGGGACTTTC | GTAGGAAAGTCCCATAAGGTC |
| --- | --- | --- |
| *Npc2 mutant* | CCAGGCAGCACGGATGTC | GCCAGGGTTTTCCCAGTCA |
| *Npc2 wild-type* | TGTGGCTCAGTGGCTTAGG | CCAGGAAGGGATTTCACACA |
